# Metabolic Connectivity Gradients of the Human Brain

**DOI:** 10.64898/2026.01.28.702441

**Authors:** Hamish A. Deery, Chris Moran, Emma X. Liang, Gary F. Egan, Sharna D. Jamadar

## Abstract

The functional architecture of the brain is organised along continuous, macro-scale gradients. However, it is unknown if the metabolic architecture of the brain displays similar gradient characteristics. Here, we use functional positron emission tomography (fPET) with 18F-fluorodeoxyglucose (FDG) to characterise the brain’s metabolic connectivity gradients and determine how neurobiological mechanisms shape these gradients to support cognition across the adult lifespan. We identified four principal metabolic connectivity gradients, with the primary axis recapitulating the canonical unimodal-to-transmodal hierarchy from other imaging modalities. Subsequent gradients delineated specialised dimensions of metabolic organisation, including association system differentiation, hemispheric asymmetry, and sensory system segregation. These gradients were coupled to cortical thickness, baseline rates of glucose metabolism, blood flow, and gene expression related to energy metabolism, such that transmodal, control and default mode poles were more metabolically active, more interconnected, had greater cortical thickness, and were more strongly related to the expression of genes related to cellular energy production, than unimodal and sensory poles. A reduction in gradient strength at the gradient poles was associated with older age and predicted worse cognitive performance. We conclude that the brain’s metabolic organisation constitutes a genetically grounded, structurally and energetically constrained gradient hierarchy that supports cognitive function and undergoes a reorganisation in ageing.

**Teaser:** The brain’s metabolic architecture forms macro-scale gradients linking genes, neurobiology, and cognitive ageing.

## Introduction

The human brain is organised to minimise energy consumption while maximising neural computation [1]. This balance is constrained by the structural and functional organisation of the brain [2]. While the organisation of the brain has long been conceptualised as a network of bounded regions, recent work has indicated that the brain is organised along a set of continuous, macro-scale gradients [3-5]. The most dominant of these gradients extends from unimodal sensory to transmodal associative cortices, and is underpinned by microscale variations in cytoarchitecture and gene expression [5-8]. This unimodal-transmodal gradient is complemented by other axes creating a multidimensional hierarchy of the brain [5, 6, 8, 9].

Functional brain gradients interact to generate state-specific connectivity patterns [10] with variability in gradient expression corresponding to distinct patterns of thought. The gradient framework guides our understanding of how information flows across networks and how macro-scale brain organisation gives rise to diverse cognitive and behavioural states [11]. For example, greater expression of transmodal regions aligns with complex cognition and self-referential thought, whereas stronger expression of visual-sensorimotor regions corresponds to problem-solving and interactions with the external environment [10, 11]. Viewed as a dynamic system, gradient-driven brain states can emerge through transient shifts in the strength of a gradient pole, fluctuations that increase functional segregation between opposing regions, or coordinated coupling between gradients [10].

Conceptualising functional brain gradients as a dynamic system also enables systematic investigation of how large-scale cortical organisation is altered in health and disease, including normative ageing. For example, alterations in brain gradients are observed in psychiatric and neurodegenerative conditions, such as Alzheimer’s disease [7, 9, 12]. While the principal gradients remain stable across the adult lifespan, their internal structure is altered in older people [13]. Specifically, transmodal networks, such as the default mode and frontoparietal systems, become more dispersed in gradient space with age, a change associated with worse cognitive performance [13, 14]. This suggests that the maintenance of a well-defined functional gradient structure is crucial for cognitive health in later life.

Much of what is known about functional gradients has been derived from functional magnetic resonance imaging (fMRI) studies. fMRI measures blood oxygenation level dependent (BOLD) signals, which are driven by the haemodynamic response to neuronal activity [15]. The temporal coherence of spatially distinct haemodynamic responses form functional brain networks and are central to the concept of the *functional connectome*. However, fMRI-derived networks are but one of a number of brain connectomes that include molecular [16], electromagnetic [17], and metabolic [18, 19] networks. With the increasing recognition of the potential causal role that cerebral metabolic dysfunction plays in neurodegenerative and psychiatric conditions [20], recent interest has focused on delineating and understanding coherent functional metabolic signals and the metabolic connectome of the brain [1, 16].

The brain’s energetic demands for neural computation are met through the vascular delivery of cerebral blood with glucose its primary fuel [21]. Measuring the haemodynamic response using fMRI does not directly measure the metabolic costs of synaptic activity, rather, it indexes a change in a mixture of blood flow, glucose metabolism, and oxygen consumption that occurs concomitantly with synaptic activity [18, 22, 23]. Recent advances in positron emission tomography (PET) now allow the measurement of metabolic connectivity by providing a direct, dynamic measure of glucose metabolism at the post synaptic neuron through the use of 18F-fluorodeoxyglucose (FDG-PET) [24-27]. Compared to fMRI-derived haemodynamic functional connectivity, metabolic connectivity is more strongly related to cognitive performance and is a better predictor of normative ageing [19, 28, 29]. The recent characterisation of cerebral ‘glucodynamics’ (time-variant cerebral glucose metabolism) from fPET has further enabled the discovery of dynamic metabolic network state transitions, which are stronger predictors of cognition than haemodynamic measures [28, 30, 31]. However, it is currently unknown if the metabolic network of the brain displays gradient characteristics. Given the recent advances in the measurement of glucodynamics and the lack of understanding of the role of metabolic connectivity in the functional architecture of the brain, our primary objective is to explore the extent to which the organisation of functional gradients reflects the brain’s metabolic architecture. On this basis, our study has four aims.

First, we aim to characterise the metabolic connectivity gradients of the human brain. Given the dominance of the canonical unimodal-transmodal gradient across brain structure, function, genetics, development and evolution [32], we hypothesise that it is also a principal feature of the brain’s metabolic organisation. Based on consistent prior evidence of a partial alignment between metabolic and functional brain networks [18, 28, 29], we hypothesise a commensurate, moderate overlap in metabolic and functional connectivity gradients. Second, we aim to elucidate how metabolic connectivity gradients are related to other features of neurobiology. We hypothesise that metabolic connectivity gradients are coupled to baseline levels of glucose metabolism, regional blood flow and glucodynamics, as well as cortical thickness and the expression of genes involved in metabolic pathways. Third, we examine age-related differences in metabolic connectivity gradients, hypothesising that older adults will exhibit a reduction in gradient strength, particularly at the gradient poles. Fourth, to link metabolic connectivity gradient characteristics to cognition, we assess the association between the gradients and cognitive task performance. We hypothesise that better cognitive performance will be associated with stronger gradient expression at the gradient poles.

## Results

We investigated metabolic connectivity gradients of the human brain using data from 84 healthy adults in the Monash *MetConn* dataset [19, 33], that included 40 younger (aged 20 to 42 years) and 44 older (66 to 86 years) people (see Supplement 1). Participants completed a simultaneous 90-minute resting-state FDG-PET and BOLD-fMRI scan, including arterial spin labelling, to measure metabolic and functional connectivity and cerebral blood flow. FDG-PET data was acquired using functional PET (fPET) [25-27]; with the bolus/infusion approach [34, 35] and 16sec frame duration. Metabolic gene expression data was sourced from the Allen Human Brain Atlas [36]. Participants also completed cognitive assessments measuring memory, processing speed, response inhibition, and task switching (see Supplement 2).

Participants’ metabolic gradients were derived from 400 x 400 metabolic connectivity matrices using diffusion map embedding [37]. To assess the neurobiological basis of these gradients, we correlated their loadings with baseline regional glucose metabolism (CMR_GLC_), cerebral blood flow, cortical thickness, metabolic network degree, functional connectivity gradient loadings and gene expression. We examined age-related differences by comparing the gradient loadings of younger and older adults. Finally, we tested the predictive value of metabolic gradients for cognition.

### Principal metabolic gradients

First, we characterised the brain’s metabolic connectivity gradients. To test our hypothesis that a canonical unimodal-transmodal gradient is a feature of the brain’s macro-scale metabolic organisation, we applied diffusion map embedding to reduce the metabolic connectivity matrices to their dominant components. The resulting gradients represent axes of the connectivity space, with regional values indicating their position along continuous dimensions that capture the principal patterns of metabolic connectivity [4]. The polarity of “positive” and “negative” gradient loadings is arbitrary and unitless, simply distinguishing whether regions oppose one another along the gradient.

Four metabolic connectivity gradients were identified, explaining 41%, 11%, 22% and 26% of the variance, respectively. The gradients are displayed on the brain surface in Figure 1a-d and in 3-D gradient space in Figures 1e-f. To characterise the correspondence of the gradients to functional network organisation, the average metabolic network loadings from the Schaefer networks [38] are also shown in Figure 1g, with each network sorted highest to lowest by gradient loading and hemisphere.

**Figure 1.**
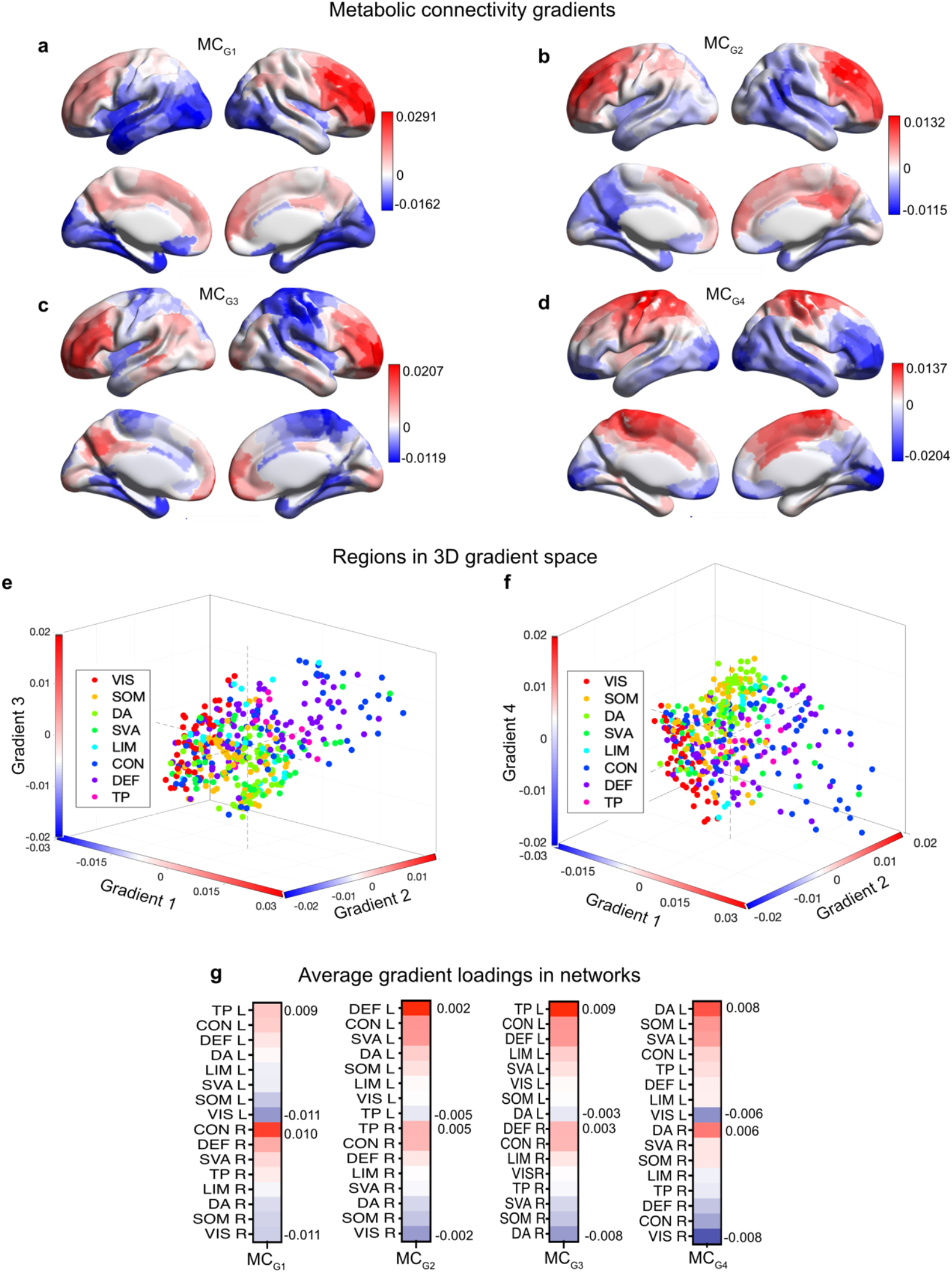
Metabolic gradients charactertistics. Whole sample (N=84) brain surface plots of four metabolic gradients (a-d). Metabolic gradients in 3D space, including gradients 1-3 (e) and 1,2 and 4 (f). Average metabolic network loadings (g), each sorted highest to lowest by gradient loading and hemisphere. VIS = Visual; SOM = somatomotor; DAN = dorsal attention; SVA = salience ventral attention; LIM = limbic; CON = control; DEF = default; TP = temporal parietal; and L = left and R = right hemisphere.

Metabolic connectivity gradient 1 (MC_G1_) represents the principal axis of the brain’s metabolic connectivity network and is characterised by a transmodal-unimodal hierarchy (Figure 1a, e and f). The positive pole was defined by transmodal areas, including prefrontal regions of the control, default, salience ventral and temporal parietal networks, as well as the precuneus and intraparietal lobule (Figure 1g). The negative pole was anchored by unimodal visual and somatomotor regions. This axis was notably stronger in the right hemisphere (Figure 1a and g), where control and default network regions showed their highest loadings. This gradient is consistent with the unimodal-to-transmodal gradient of the brain [5] and reflects a hierarchy from higher-order cognition and self-generated thought through to sensory perception.

The second metabolic connectivity gradient (MC_G2_) separated hemisphere lateralised networks (Figure 1b). The regions of the temporal parietal network showed complete hemisphere lateralisation, with positive gradient loadings for the right hemisphere regions and negative loadings for the left. In the left hemisphere, the positive pole had loadings from mostly control and default regions (Figure 1g). The right hemisphere also had positive loadings from regions of the control, default and temporal parietal networks, which were strongly opposed to sensorimotor and visual regions at the negative pole. This pattern suggests that MC_G2_ reflects hemispheric differentiation of associative systems. In other words, while MC_G1_ orders regions by their level of abstraction (from cognition to sensation), MC_G2_ arranges them by their functional affiliation and hemispheric dominance.

Metabolic connectivity gradient 3 (MC_G3_) reflects a hierarchy from cognitive control through to visuospatial attention regions (Figure 1c). The positive pole included regions of the dorsolateral prefrontal cortex, inferior parietal lobule and left temporal regions (Figure 1g). In contrast, the negative pole was dominated by dorsal attention and visual networks in the right hemisphere, including regions of the frontal eye fields and intraparietal sulcus. This gradient captures a hierarchy of cognitive control and spatially-focused attention.

Metabolic connectivity gradient 4 (MC_G4_) revealed a visual-cognitive to affective-temporal integration axis, characterised by distinct network configurations at the poles (Figure 1d, g). The negative pole comprised a combination of visual processing regions with cognitive control systems (prefrontal regions) particularly strong in the right hemisphere and suggesting a network for perceptually-guided executive function. Conversely, the positive pole was anchored by attention and somatomotor networks, including frontolimbic and medial temporal regions (default parahippocampal gyrus, limbic temporal pole, salience prefrontal cortex) involved in affective processing and memory integration. This organisation highlights an axis of externally-oriented perceptual-cognitive integration and internally-oriented emotional and memory processing, representing a distinct hierarchy that bridges sensory and cognitive domains.

Taken together, these four gradients reveal a multi-dimensional metabolic connectivity architecture anchored by a primary unimodal-transmodal hierarchy (MC_G1_). This foundational axis is complemented by gradients of hemispheric specialisation (MC_G2_), executive-control versus visuospatial attention (MC_G3_), and an axis segregating perceptual-cognitive from affective-temporal processing (MC_G4_). These results demonstrate that the brain’s metabolic connectome is defined by intersecting principles of hierarchical processing, lateralisation, and functional specialisation.

### Metabolic and functional gradients

Next we examined the relationship between the four metabolic connectivity gradients and fMRI-based functional connectivity gradients. We first describe the fMRI gradients, derived from diffusion map embedding of 400 x 400 functional connectivity matrices. We then use correlation analyses to test our hypothesis that metabolic connectivity gradients will show moderate overlap with functional gradients.

Four functional connectivity (FC) gradients were identified, accounting for 37%, 20%, 32%, and 12% of the variance, and together delineating multiple axes of the brain’s functional organisation (Figure 2). The FC gradients are described in detail in Supplement 3. Briefly, the principal functional connectivity gradient (FC_G1_) reflected the core transmodal-to-unimodal axis, spanning from default and control networks to visual and somatomotor regions (Figure 2a). FC_G2_ captured a distinction between an externally-oriented dorsal attention and somatomotor system and internally oriented transmodal regions (Figure 2b). FC_G3_ differentiated the brain’s intrinsic, self-referential system (default and temporal-parietal networks) from primary sensorimotor cortices (Figure 2c). FC_G4_ revealed a fundamental sensory segregation, distinguishing somatomotor from exteroceptive visual perception (Figure 2d).

**Figure 2.**
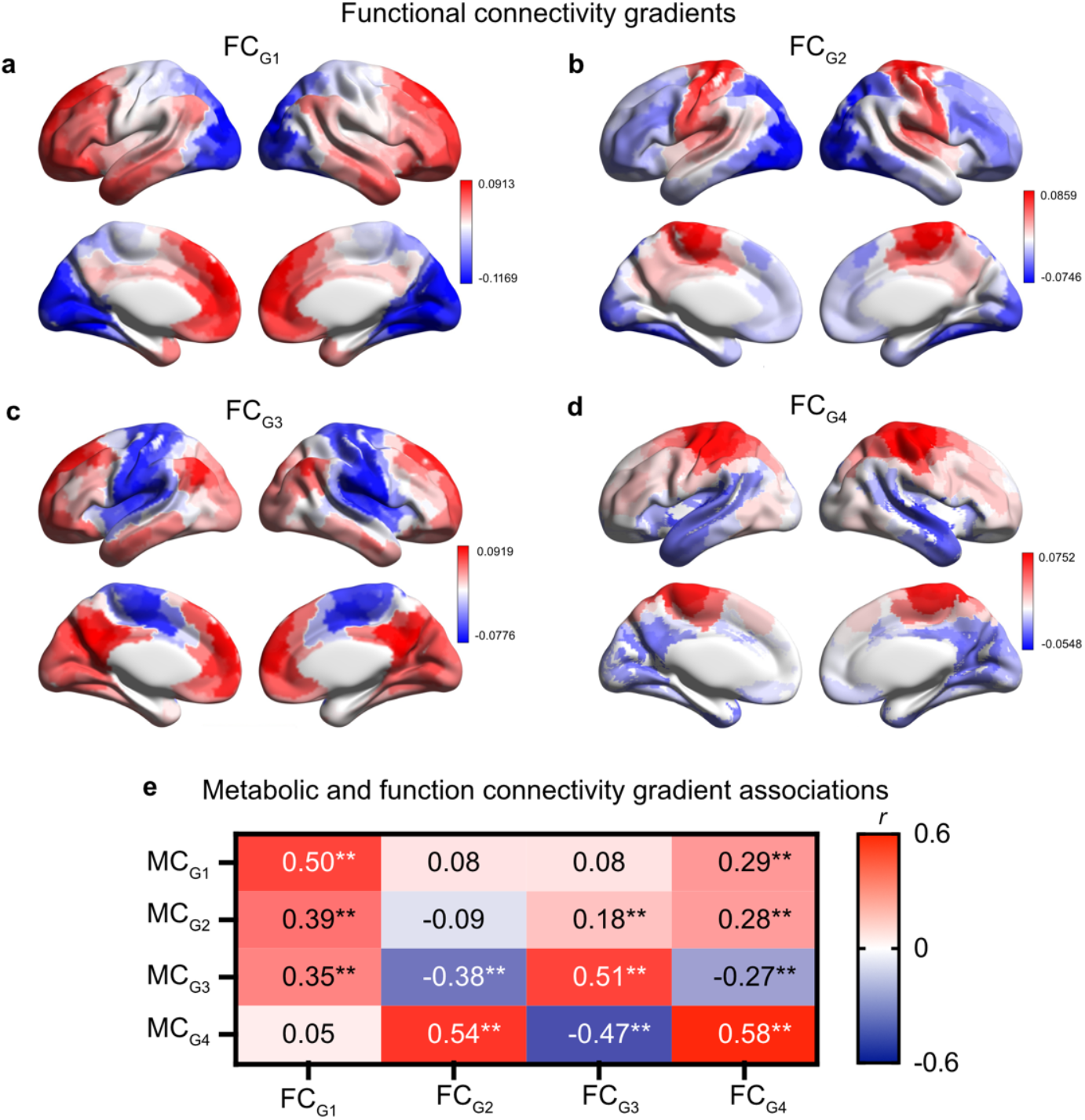
Functional connectivity gradients and their assocition with metabolic connectivity gradients. Surface plots of four functional connectivity gradients (a-d) and correlation of the four functional and metabolic connectivity gradient loadings (e). *p-FDR < 0.05; **p-FDR < .001.

Qualitatively, MC_G1_, MC_G3_, and MC_G4_ resembled their counterparts in the FC data, but MC_G2_ and FC_G2_ appeared to differ in their spatial topology. Gradients 1, 3, and 4 accounted for the highest level of variance in each modality. MC_G1_, MC_G3_ and MC_G4_ showed significant positive correlations with their corresponding functional gradients (r = 0.58 to 0.50, p-FDR < .001; Figure 2e). MC_G3_ was uniquely characterised by its negative correlation with FC_G2_ and FC_G4_ (r = -0.38 and r = -0.27, p-FDR < .001). Higher metabolic connectivity in networks for sensory attention coincided with lower functional connectivity in executive networks, and vice versa. For example, higher loadings in regions at the positive pole of MC_G3_ (dorsolateral prefrontal cortex, inferior parietal lobule and left temporal regions) corresponded to lower loadings at the positive pole of FC_G2_ (dorsal attention regions) and FC_G4_ (somatomotor and dorsal attention regions). Similarly, higher loadings in regions at the positive pole of MC_G4_ (dorsal attention and somatomotor regions) corresponded to lower loadings at the positive pole of FC_G3_ (default mode and temporal parietal networks).

Together, these results indicate moderate alignment between the brain’s primary metabolic and functional connectivity gradients, with some key exceptions in specific functional systems. The moderate strength of these relationships is informative. The organisation of metabolic connectivity reflects the underlying metabolism that supports, but is not perfectly mirrored by, functional connectivity. While the BOLD signal is an indirect measure of neuronal activity, FDG-fPET provides a more direct measure of metabolism and reflects the fundamental metabolic substrate of neuronal activity [24]. Our results add to a growing body of evidence that glucodynamic measures provide important new information about brain function [18, 28, 29] and their divergence from fMRI measures reinforces the value of fPET to our understanding of brain organisation.

### Metabolic connectivity gradient and brain physiology

Next, we examined the relationship between the four metabolic connectivity gradients and measures of neurobiological properties that shape metabolism and function network organisation in the brain. We used correlation analyses to test our hypothesis that metabolic gradients are associated with baseline rates of glucose metabolism, blood flow, glucodynamics and cortical thickness. We first describe the common physiological associations across the four metabolic gradients. We then describe the unique gradient-specific neurobiological correlates.

#### Common physiological associations across gradients

The neurobiological properties are shown on the brain surface in Figure 3a-f. The four metabolic connectivity gradients demonstrated significant associations with the neurobiological measures, establishing their broad physiological bases (Figure 3g). The first three gradients were positively associated with CMR_GLC_, (r = 0.39 to 0.21, p-FDR < 0.011), fPET metabolic network degree (r = 0.76 to 0.63, p-FDR < .001) and cortical thickness (r = 0.29 to 0.23, p-FDR < .001). In other words, the positive poles (i.e., transmodal and executive function poles of MC_G1_ and MC_G3_, and right-hemisphere control and default mode pole of MC_G2_), were more metabolically active, more interconnected, and had greater cortical thickness than their opposing negative poles (mostly unimodal or sensory regions). Each metabolic gradient was also positively associated with cerebral blood flow (r = 0.29 to 0.10), including significant associations for MC_G1_, MC_G2_ and MC_G4_, indicating higher perfusion at the transmodal than unimodal gradient poles. Together, these shared associations confirm that the brain’s metabolic organisation is systematically related to its baseline rates of perfusion and glucose metabolism, cortical thickness and metabolic network organisation.

**Figure 3.**
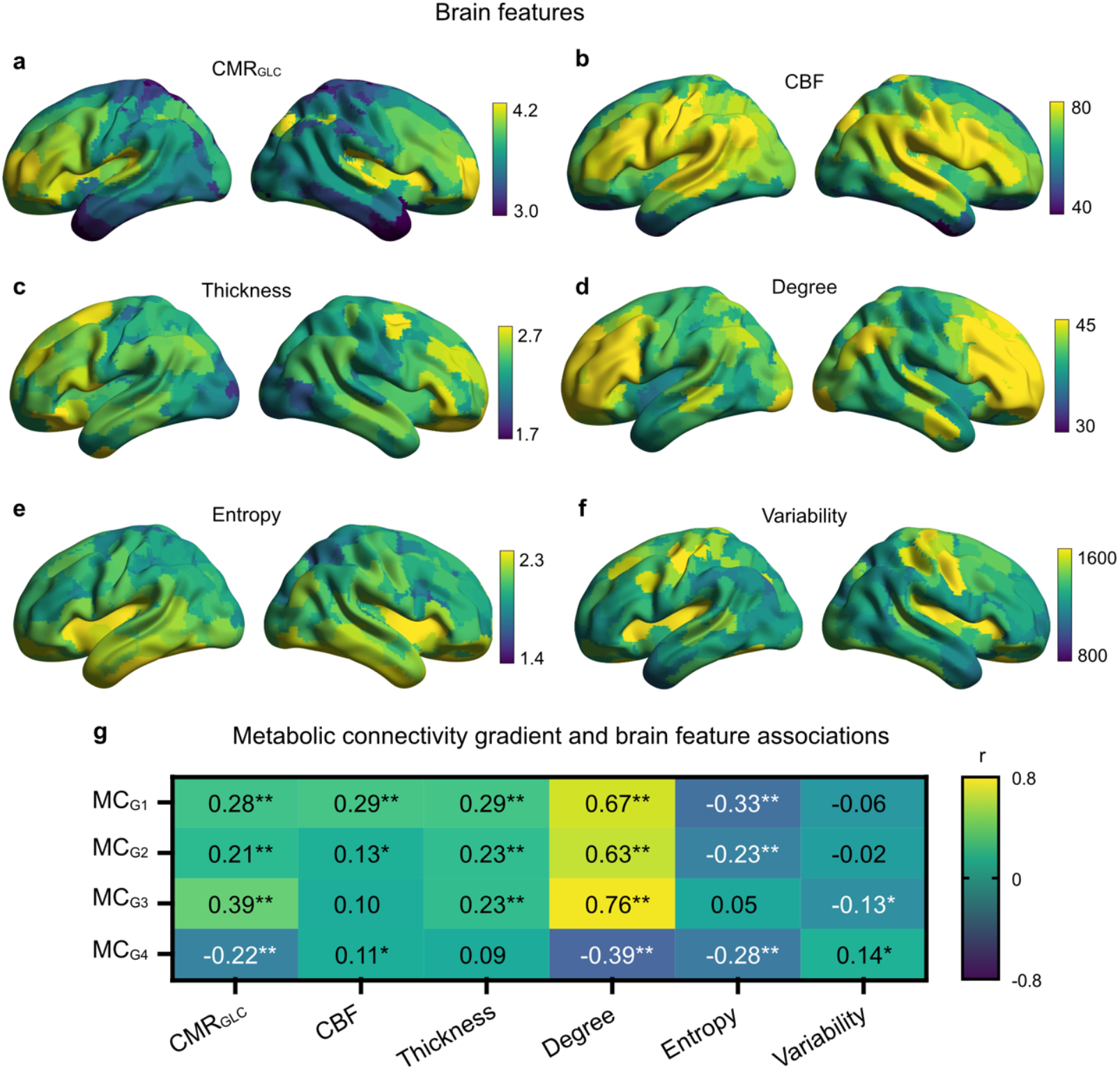
Metabolic connectivity gradients and neurobiological brain feature associations. Surface plots of neurobiologial brain measures, including regional cerebal metabolic rate of glucose (CMR_GLC_) (a), cerebral blood flow (CBF) (b), cortical thickness (c), fPET degree (d), fPET entopy (e) and fPET variabiloty (SD) (f). Correlation of metabolic connectivity gradients loadings with the neurobiologocal brain features (g). *p-FDR < 0.05; **p-FDR < .001.

#### Unique gradient-specific correlates

Beyond these common relationships, metabolic connectivity gradients MC_G3_ and MC_G4_ exhibited a distinct physiological signature. MC_G4_ showed a unique negative associated with CMR_GLC_ (r = -0.22, p-FDR < .001) and fPET metabolic network degree (r = -0.39, p-FDR < .001) (Figure 3g). These negative associations indicate an inversion of the relationship between physiology and the gradient, such that the positive pole (dorsal attention and somatomotor networks, including auditory regions) had lower cerebral perfusion and baseline metabolism than the opposing pole (visual regions).

We have recently reported that the variability and complexity of the fPET timeseries, which we have called ‘glucodynamics’, is an important feature of the brain’s energetic architecture, and is highly predictive of age and cognition, more so than classic measures of CMR_GLC_ and fMRI-derived metrics [30, 31]. Glucodynamic measures showed negative associations with the metabolic gradients. Higher entropy (complexity) of the fPET timeseries was significantly associated with lower MC_G1_, MC_G2_ and MC_G4_ expression (r = -0.33 to -0.23, p-FDR < .001). Higher variability of the fPET timeseries was significantly associated with lower gradient expression (r = -0.13, p-FDR < .05). For example, for MC_G1_, the positive (prefrontal regions of the control, default, salience ventral and temporal parietal networks) had lower entropy and variability than the negative pole (visual and somatomotor regions). Together, these findings indicate systematic network-level differences in dynamic glucose utilisation that map onto the brain’s metabolic hierarchy. Transmodal and executive regions exhibit lower glucodynamic complexity, consistent with flexible state transitions emerging from a relatively stable metabolic baseline. In contrast, unimodal regions show greater glucodynamic variability, potentially reflecting heightened metabolic responsiveness to fluctuating sensory demands.

Taken together, our findings reveal that the metabolic connectivity network organisation of the brain follows conserved hierarchical principles, where transmodal regions consistently receive greater physiological resources, and express higher metabolic and functional network connectivity. There are also specialised exceptions, where particular functional systems like the dorsal attention network operate with distinct neurobiological signatures (lower CMR_GLC_ and perfusion), and sensory-motor and attention system demonstrate more complex time-variant glucose use.

### Functional connectivity gradient and brain physiology

Although not a primary focus of the current study, we also examined the relationship between the four fMRI-derived functional connectivity gradients and the neurobiological brain features, comparing the results to those found for the metabolic connectivity gradients. The results are reported in Supplement 4. Briefly, the functional connectivity gradients were less consistently associated with CMR_GLC_, cerebral blood flow and cortical thickness and expressed different patterns of network connectivity degree compared to the metabolic connectivity gradients. In contrast, glucodynamic and haemodynamic complexity and variability consistently mapped onto their respective gradient hierarchies. Together, these results further support our hypothesis that metabolic and functional connectivity gradients show moderate similarity and are differentially associated with other neurobiological brain features.

### Gene expression

Next we examined the multivariate associations between the metabolic connectivity gradients and gene expression using partial least square correlation analysis. Four statistically significant latent components were found (Figure 4a-d), each with significant positive correlations, indicating that higher gene expression is associated with higher metabolic gradient loadings. The first component (LV1) was the most dominant, explaining 61% of the covariance and with a significant correlation of r = 0.46 (p < .001). The second component (LV2) accounted for a further 36% of the covariance, also with a correlation of r = 0.46 (p < .001). While two subsequent components (LV3 and LV4) explained smaller proportions of variance (2% and 1%, respectively), their correlations were also significant (r = 0.29, p < .001 and r = 0.35, p < .001).

**Figure 4.**
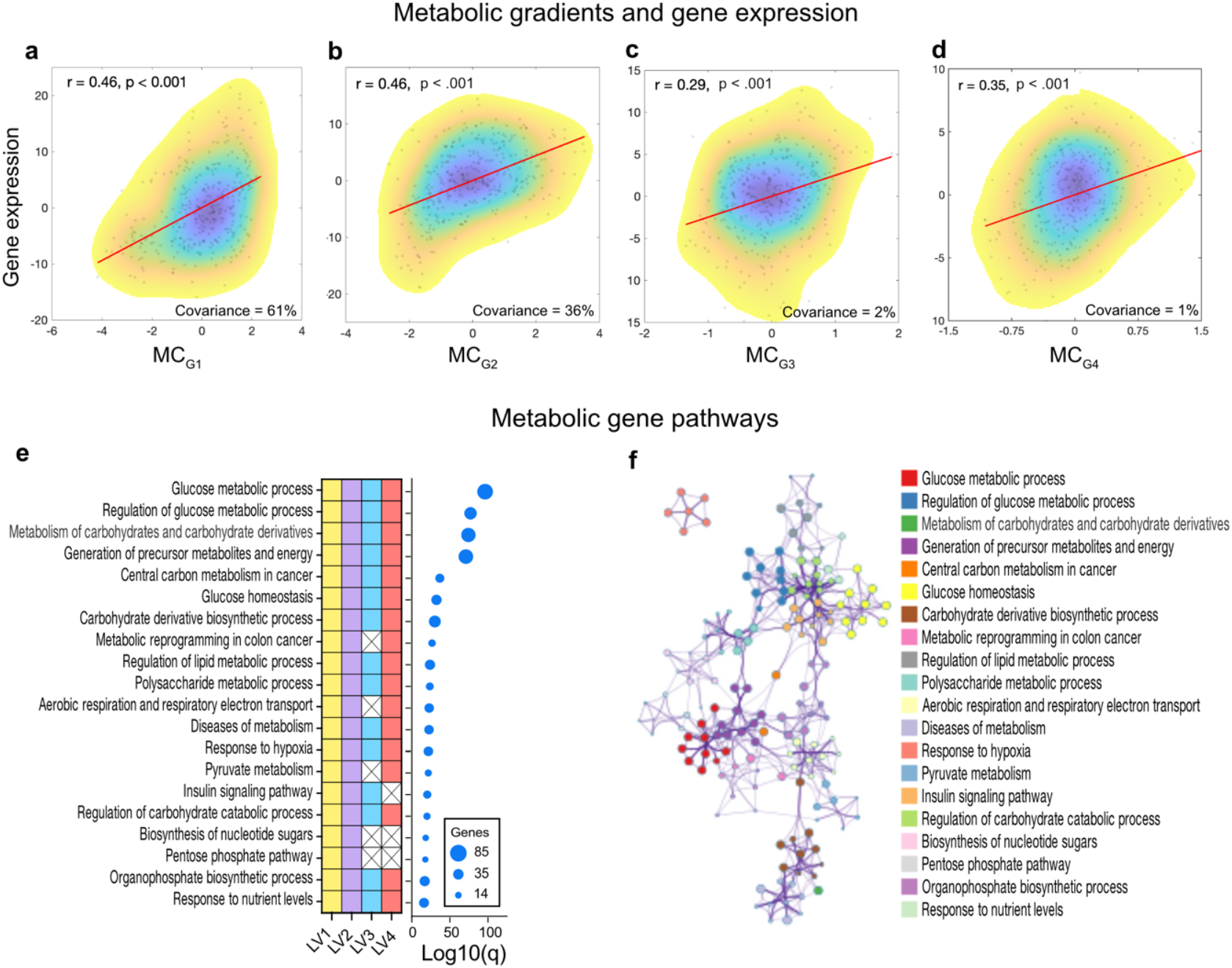
Metabolic gradients and gene expression associations. Correlations between the four metabolic gradients and 447 gene expression data from partial least square correlation analysis in the whole sample (N = 84) (a-d). The correlation coefficient (r) and p-value quantify the strength and significance of the association. The significant unique genes from LV1 (n = 150), LV2 (130), LV3 (36) and LV4 (52) were used in gene enrichment analysis (e). Top 20 clusters are shown with their representative enriched terms the number of genes in the given ontology term. q-values are calculated using FDR-correction and are shown as in log base 10 (Log10(q)). Enrichment terms with a p-value < 0.01, a minimum count of 3, and an enrichment factor > 1.5 (the enrichment factor is the ratio between the observed counts and the counts expected by chance) were grouped into clusters based on their membership similarities (f). Terms with the best p-values from each of the 20 clusters were selected and those with a similarity > 0.3 are shown with connected by edges. Gene enrichment analyses was undertaken at metascape.org [39].

Bootstrap stability analysis revealed multiple robust gradient-gene expression associations, with LV1 exhibiting the strongest effect, contributing 190 significant genes (150 unique), followed by LV2 (152 genes, 120 unique), LV4 (68 genes, 52 unique), and LV3 (45 genes, 36 unique). Functional enrichment analysis of this gene set [39] revealed a highly significant representation of biological pathways and processes related to energy metabolism (Figure 4e-f). The most significantly enriched terms were directly associated with glucose metabolism, including “glucose metabolic process” (85 genes, Log10(q) = -96.3), “regulation of glucose metabolic process” (54 genes, Log10(q) = -77.1), “metabolism of carbohydrates and carbohydrate derivatives” (71 genes, Log10(q) = -74.3), and “generation of precursor metabolites and energy derivatives” (75 genes, Log10(q) = -71). These processes were complemented by strong enrichment in broader metabolism and key regulatory pathways, such as insulin and glucagon signalling, glucose homeostasis and lipid metabolic regulation. Collectively, these results indicate that the strength of metabolic connectivity gradients of the brain is tied to the expression of a functionally coherent gene set of cellular energy production and metabolic function.

### Age group differences in metabolic gradients

Next we tested for age group differences in the gradient measures. To test our hypothesis that older adults would have lower gradient strength than younger adults, particularly at the gradient poles, we used independent sample *t*-tests. The magnitude measures at the poles were lower in the older group (Figure 5a-b) than the younger group. Among the top 10% of positive-loading regions in the whole sample, older adults showed significantly lower strength of loadings than younger adults (t = 2.80, p = 0.007). A similar pattern was observed for the 10% most negative loadings in the sample, where older adults also showed lower strength of loadings, although this difference did not reach significance (t = 1.72, p = 0.090). Older adults tended to exhibit a lower sum of absolute gradient loadings (t = 1.50, p = 0.142) and lower standard deviation of gradient loadings (t = 1.60, p = 0.116), although both differences were not significant (Figure 5c-d). These results indicate a pattern of modestly decreased strength of gradients in older than younger adults, primarily at the positive or transmodal poles of the gradients.

**Figure 5.**
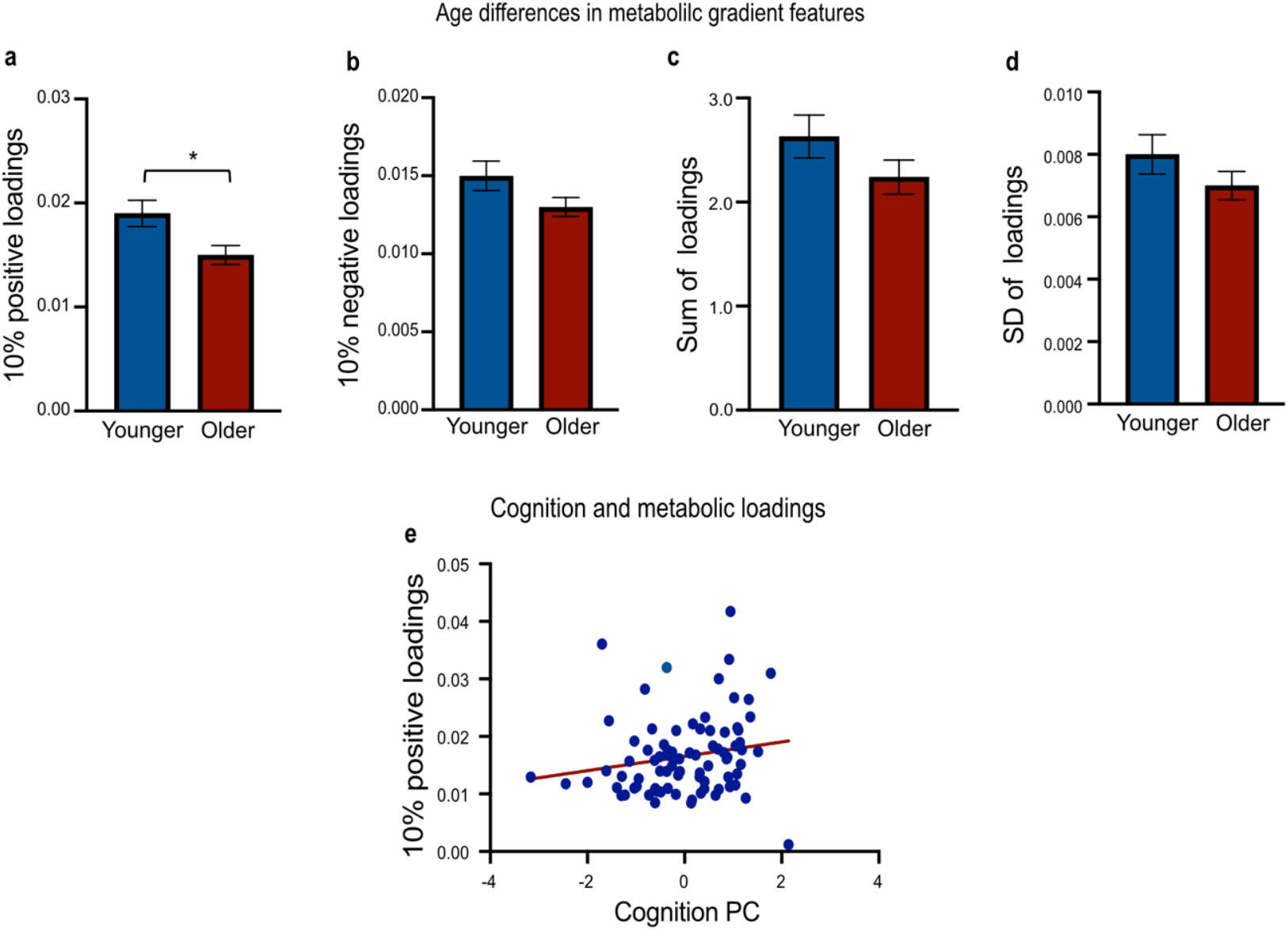
Age group differences in metabolic gradients features and their association with cognition. Younger (N = 40) and older (N = 44) adult mean (± SE) top 10% of positive (a), and negative (b) gradient loading regions, and sum (c) and SD (d) of all absolute gradient loadings. Association between top 10% of positive-loading regions of the metabolic connectivity gradients and cognition (e). Cognition PC is the principal component score based on Z-scored performance for HVLT delayed recall, reaction time in the switching and a stop signal tasks, and the number of correct responses in the digit substitution task. Reaction time measures were multiplied by -1 so that higher scores correspond to better performance. In multiple regression analysis of the metabolic gradient features and cognition PC, stronger metabolic gradient loadings among the top 10% of positive-loading regions was associated with better cognitive performance (ß = 0.59, p = 0.012). *p < .05.

### Metabolic gradients and cognition

Lastly, we assessed the relationship between metabolic gradients and cognition. To test our hypothesis that better cognitive performance would be associated with stronger gradient expression, particularly at the gradient poles, we used principal component analysis to reduce the dimensionality of the cognitive data and regression analyses to test its association with the metabolic gradients. Principal component analysis of the four cognitive measures identified one significant component explaining 48.9% of the variance. This principal cognitive component loaded most strongly on verbal learning (HVLT, 0.84), followed by psychomotor speed (digit symbol substitution, 0.74), inhibitory control (stop signal, 0.63), and cognitive flexibility (category switch, 0.56).

We found a statistically significant association between the gradient measures and the cognition principal component on regression analysis (F = 9.48, p = .046, R^2^ = 11.6%). Stronger metabolic gradient loadings among the top 10% of positive-loading regions was associated with better cognitive performance (ß = 0.59, p = 0.012; Figure 5e). There was a trend towards the 10% most negative loadings being associated with worse cognition, but this was not significant (ß = -0.31, p =0.110). The sum and the standard deviation of all gradient loadings were not significantly associated with cognition (ß = -2.2, p = 0.267 and ß=1.93, p = 0.332)

## Discussion

Our results establish for the first time a multi-dimensional metabolic architecture of the human brain characterised by four metabolic connectivity gradients. As hypothesised, the primary metabolic gradient recapitulates the well-established unimodal-to-transmodal hierarchy of the brain reported in other imaging modalities [5], and confirms its foundational role in brain organisation. Crucially, we demonstrate that these metabolic connectivity gradients are related to the brain’s underlying cortical thickness, baseline rates of glucose metabolism, cerebral blood flow, and genetic expression related to energy metabolism. The metabolic connectivity gradients were moderately related to functional connectivity gradients. A loss of gradient strength occurs in ageing at the positive gradient poles, which is also associated with worse cognitive performance. Taken together, our findings establish a metabolic connectivity hierarchy of the human brain that is genetically grounded, connected to cortical structure and energy supply and is predictive of ageing and cognitive performance. Our results add to the growing body of research showing the value of incorporating dynamic metabolic imaging into studies of large-scale brain organisation [19, 29, 31, 33, 40]. fPET offers a unique window into the energetic constraints shaping neural communication, particularly in contexts such as ageing and neurodegenerative disease where vascular and metabolic processes are differentially affected [41].

This primary metabolic connectivity gradient (MC_G1_) identifies a segregation between processes that support sensory–motor interactions with the external environment and those that enable complex cognition and memory-driven actions [11]. Subsequent gradients revealed specialised dimensions of metabolic organisation, including association system differentiation, hemispheric asymmetries, and sensory system segregation. MC_G2_ and MC_G3_ demonstrate that hemispheric specialisation is a core component of the brain’s metabolic connectivity hierarchy. These results provide a metabolic basis for established neuropsychological models [42] and build on fMRI studies of lateralisation of attention [43, 44] and other networks supporting cognitive performance [45]. The presence of these asymmetries in metabolic connectivity gradients suggests that the brain’s functional lateralisation, which has been connected to cognitive efficiency, is supported by an underlying metabolic organisation that favours specialised, energy-efficient unilateral processing over more costly bilateral integration [46].

MC_G4_ reveals a fundamental, bilateral hierarchy dedicated to sensory integration, spanning from visual perception to somatomotor and auditory processing. MC_G4_ was moderately correlated with a similar functional connectivity gradient and was characterised by regions that integrate multisensory information [47-49]. The presence of this gradient indicates that the brain invests metabolic resources in the energetically costly but crucial integration of sensory and attentional processes that underlie adaptive behaviour.

The metabolic connectivity gradients also reflect a conserved neurophysiological hierarchy in which the brain’s most metabolically demanding regions at the transmodal poles receive a preferential allocation of energetic resources to support higher order cognitive processes. There is a convergence of high baseline cerebral blood flow and glucose metabolism in those regions, aligning with the ‘metabolic cost’ hypothesis [50, 51]. This convergence suggests that the brain’s functional hierarchy is fundamentally constrained by its vascular and metabolic systems [21], where the high energetic cost of complex, integrative computation, particularly in frontoparietal regions, is supported by an adaptive vascular and metabolic system [1, 52]. Importantly, the positioning of transmodal regions at the poles of metabolic connectivity gradients likely reflects their role as metabolically costly, integrative hubs with high baseline energetic demands and widespread connections across the brain that support effective information transfer [19, 32].

We found that transmodal and executive regions of the metabolic gradients also exhibited greater cortical thickness than unimodal regions, suggesting coupling between macro-scale metabolic connectivity and underlying cytoarchitecture. Our results align with previous work showing that cortical thickness is systematically related to the structural hierarchical organisation of the cortex [53] and that cortical thickness and synaptic loss in ageing and early Alzheimer’s disease (AD) are major structural correlates of cognitive impairment [54]. There is also an anterior-to-posterior gradient in the density of neurons that is believed to foster the integration of information from sensory areas to the frontal cortex [55]. From this perspective, metabolic connectivity gradient organisation may be coupled to synaptic density, laminar structure and broader cellular and circuit-level architecture that underpins complex information integration [56]. Our results further support the view that the brain’s functional hierarchy is fundamentally constrained by metabolic resources, with transmodal frontoparietal regions representing metabolically costly but computationally flexible nodes within the cortical hierarchy (also see [57]). Recent advances in PET imaging that allow for a direct measure of synaptic density in the human brain *in vivo* [58] could be used to directly test the relationship between metabolic gradients and synaptic density in the future.

Our results suggest a mechanistic link between macro-scale metabolic gradient organisation and gene expression. Four latent variables were enriched for 338 unique genes involved in glucose metabolism, metabolic pathways, and insulin and glucagon signalling. As hypothesised, this finding suggests that the spatial hierarchy of the brain’s metabolic connectivity gradients is genetically encoded, shaped by regional variation in the expression of genes that govern cellular energy production and regulation. In other words, the architecture of the brain’s macro-scale functional and metabolic connectivity is constrained by the molecular machinery that fuels its functional capacities [1]. Converging evidence indicates that transmodal regions and their metabolically demanding hub areas arise from coordinated transcriptional programs that are shaped by physical and geometric constraints on cortical organisation, early patterning processes anchored in primary sensory fields, and later self-organising, activity-dependent mechanisms (see [32] for review). By directly linking gene expression to the macro-scale metabolic organisation of the brain, our results support earlier work that shows gene transcription underlies structural and metabolic brain alterations and mediates risk for cognitive changes and psychiatric and neurodegenerative disease [59-63].

Here, we reveal that regions with higher variability and complexity of dynamic glucose use are at the unimodal poles of the metabolic connectivity gradients, such as regions of the sensory and attention systems. Greater variability in neuronal firing patterns is believed to represent greater system adaptability and hence the information processing capacity of the brain [64-68]. We recently reported that higher variability and complexity of glucodynamic signals across the brain enable a richer repertoire of network state transitions over time [31, 69]. Together these results suggest that there are differences across networks in terms of their dynamic glucose use and their position on the metabolic hierarchy. Transmodal and executive regions show lower glucodynamic complexity, suggesting an ability to shift between network states from a relatively stable baseline of glucose metabolism. Conversely, higher glucodynamic variability in unimodal regions appears to reflect a readiness to respond to changing sensory information, which may make them more responsive to changes in the sensory environment. A breakdown in this balance could be implicated in neurodegenerative or psychiatric conditions. For example, excessive rigidity or complexity in glucose dynamics and an altered metabolic hierarchy in specific functional networks may drive the cognitive changes seen in dementias, such as Alzheimer’s [70].

Our finding that older adults had weaker metabolic connectivity gradients at the positive poles and that this corresponds to regions and networks that support higher order cognitive processes builds on the previously reported functional and structural gradient alterations in ageing [13, 14]. Ageing, which is classically associated with reductions in cortical thickness, cerebral glucose metabolism, and cerebral perfusion [71], is also associated with reductions in metabolic connectivity and glucodynamic variability and complexity [19, 30]. Our results indicate that the cognitive decline associated with ageing, in which metabolic disturbance is thought to be a causal factor [20], may be driven by changes at the metabolic gradient poles. Enhanced age-related decline at the positive metabolic gradient poles may explain why the earliest changes in ageing are seen in cognitive functions that rely on executive, control, and default mode networks [72, 73]. In addition, the results indicate there is a metabolic basis for age-related changes in the functional hierarchical organisation of the brain that is not captured by fMRI measures of haemodynamic connectivity alone. Further work is required to understand the implications of the compression of the hierarchical organisation we report and the role of maintaining the integrity of metabolic architecture in cognitive ageing.

As hypothesised, we observed a moderate association between metabolic and functional connectivity gradients, indicating partial convergence but also meaningful differences between modalities. Metabolic connectivity gradients were more consistently associated with regional CMR_GLC_ and cerebral blood flow than functional connectivity gradients, suggesting that distinct biological constraints shape metabolic and functional gradient organisation. In contrast, glucodynamic and haemodynamic complexity and variability mapped more consistently onto their respective gradient hierarchies, indicating that these measures capture coordinated dynamics and systems-level network organisation. These observed differences in gradient organisation and their neurophysiological correlates further reinforce the view that dynamic metabolic and functional imaging capture complementary, non-redundant aspects of brain organisation [18, 19, 28, 29, 31, 69]. In particular, fMRI and fPET are sensitive to different facets of neuronal activity. The BOLD fMRI signal reflects a complex, indirect haemodynamic response driven by changes in cerebral blood flow, cerebral metabolic rate of oxygen, lactate production, and oxygen extraction fraction [74], and is therefore influenced by neurovascular coupling and vascular variability. In contrast, FDG-PET provides a measure more closely coupled to regional glucose uptake, reflecting cellular energy consumption that is tightly linked to excitatory synaptic activity [1, 26, 35]. Although glucose uptake is not entirely independent of perfusion, FDG-PET is less sensitive to transient haemodynamic fluctuations and offers a more direct index of the metabolic demands associated with sustained neural processing [1].

The results of this study should be considered in light of its strengths and limitations. A key strength is the use of recently developed fPET to characterise metabolic connectivity gradients. Another strength is the simultaneously acquired neuroimaging measures, allowing the association between metabolic and functional connectivity gradients and other neurobiological features to be tested. The comparison of metabolic connectivity gradients in older and younger adults is also a strength. The study’s limitations should also be considered. Because the age-related comparisons are cross-sectional, longitudinal studies are required to confirm that the observed age differences in metabolic gradients represent true within-individual changes during the course of ageing. In addition, although statistically robust, our neurobiological associations are correlational and the gene expression data based on external samples, hence they cannot establish causality.

On the other hand, the methods used here and findings also open opportunities for future research. A primary question is whether specific neuropsychiatric and neurodegenerative diseases exhibit distinct patterns or changes along metabolic gradient axes [75]. This is particularly important given that drug discovery work currently focuses on cerebral metabolic disturbance as a putative causal factor in psychiatric and neurodegenerative disease [20]. Such work could establish metabolic gradients as prognostic biomarkers for clinical staging. Longitudinal and lifespan studies could identify factors that promote accelerated changes or resilience in health and disease. Furthermore, metabolic gradients could be compared to the topology of major neurotransmitter systems (e.g., dopamine, serotonin) to establish connections between cerebral energy use and neuromodulation. Finally, combining our transcriptomic findings with proteomic and genetic data could extend beyond correlation to build causal models of how genetic variation shapes the brain’s metabolic organisation. Together, these directions position metabolic connectivity gradients as a framework for understanding the biological basis of cognition and its vulnerability in disease.

In conclusion, here we delineate a multi-dimensional architecture of metabolic connectivity gradients in the human brain. This architecture is built on a foundational unimodal-transmodal hierarchy, which is complemented by specialised axes of hemispheric specialisation for cognitive systems and a hierarchy for multisensory integration. This architecture is shaped by a conserved metabolic, vascular and structural coupling and is genetically encoded for energy metabolism. The integrity of this metabolic architecture is important for cognitive health. Attenuation of gradient strength at the transmodal poles occurs in older adults and is linked to worse cognitive performance, and provides a metabolic basis for the functional alterations observed in ageing. Furthermore, the moderate alignment of our metabolic and functional connectivity gradients reinforces the value of glucodynamics to our understanding of brain organisation and as a complement to other imaging modalities. Our findings position metabolic gradients as a central organisation principle of the brain and provide a metabolic basis for understanding cognitive vulnerabilities in ageing and disease, opening new avenues for biomarker development and therapeutic targeting.

## Materials & Methods

### Ethical Considerations

The study protocol was reviewed and approved by the Monash University Human Research Ethics Committee. Administration of ionizing radiation was approved by the Monash Health Principal Medical Physicist, following the Australian Radiation Protection and Nuclear Safety Agency Code of Practice (2005). For adult participants, a 5 mSv annual radiation exposure limit applied. The effective dose in our study was 4.9 mSv. Participants provided informed consent to participate.

### Participants

Local advertising was used to recruit 90 participants from the general community. An initial screening interview ensured that participants had the capacity to provide informed consent. Exclusion criteria were a diagnosis of diabetes, neurological or psychiatric illness or dementia. Participants were also screened for claustrophobia, non-MR compatible implants, and a PET scan in the past 12 months, and for women current or suspected pregnancy. Participants received a $100 voucher for completing the study.

Six participants were excluded from further analyses due to excessive head motion (n=2), incomplete PET scan (n=2) or being > 3 SDs in the connectivity matrix used for the metabolic gradient analysis (N = 2). The final sample included 84 participants (see Supplement Table S1).

### Demographic and Cognitive Variables

Prior to the scan, participants completed a demographic questionnaire and a battery of cognitive tests, comprised of domains validated in ageing research [76] (see Supplement for details): delayed recall from the Hopkins Verbal Learning Test (HVLT); reaction time in a task switching test to index cognitive control; reaction time in a stop signal task to measure response inhibition; and the number of correct responses in a digit substitution task to measure visuospatial performance.

### MR/PET Data Acquisition

Participants underwent a 90-minute simultaneous MR/PET scan in a Siemens Biograph 3-Tesla molecular MR scanner. They were requested to consume a high-protein and low-sugar diet for the 24 hours prior to the scan, fast for six hours and to drink 2–6 glasses of water. Participants were cannulated in the vein in each forearm and a 10ml baseline blood sample was taken. At the beginning of the scan, half of the 260 MBq FDG tracer was administered via the left forearm as a bolus. The remaining 130 MBq of the FDG was infused at a rate of 36ml/hour over 50 minutes. This combined bolus and constant infusion protocol maximises the signal-to-noise ratio over the period of the scan [40].

Participants were positioned supine in the scanner bore with their head in a 32-channel radiofrequency coil. The scan sequence started with non-functional MRI scans during the first 12 minutes, including a T1 3DMPRAGE (TA = 3.49 min, TR = 1,640ms, TE = 234ms, flip angle = 8°, field of view = 256 × 256 mm^2^, voxel size = 1.0 × 1.0 × 1.0 mm^3^, 176 slices, sagittal acquisition) and T2 FLAIR (TA = 5.52 min, TR = 5,000ms, TE = 396ms, field of view = 250 × 250 mm^2^, voxel size = .5 × .5 × 1 mm^3^, 160 slices). Thirteen minutes into the scan, list-mode PET (voxel size = 2.3 x 2.3 x 5.0mm^3^) sequences were started.

A 40-minute resting-state scan was acquired while participants watched a movie of a drone flying over the Hawaii Islands. At 53 minutes, a 5-delay pseudo-continuous arterial spin labelling (pcASL) scan was initiated (TR = 4,220 ms; TE = 45.46 ms; FOV = 240 mm; slice thickness = 3 mm; voxel size 2.5 x 2.5 x 3.0 mm^3^). Post labelling delays were 0.5, 1.0, 1.5, 2.0, and 2.5s and duration of the labelling pulse was 1.51s.

Plasma radioactivity levels were measured during the scan from 5ml blood samples taken from the right forearm, every 10-minutes, for a total of nine samples. The samples were spun in a centrifuge at 2,000 rpm for 5 minutes. 1,000-μL of plasma was placed in a well counter for four minutes and the count start time, total number of counts, and counts per minute were recorded.

### Imaging data pre-processing and brain feature extraction

The data pre-processing and brain feature measures extracted from the imaging data have been described extensively in our previous reports [28, 33, 69]. We describe them here briefly.

#### Cortical Thickness

Cortical thickness was extracted for the 400 regions of the Schaefer functional parcellation [38]. We used FreeSurfer’s *recon-all* pipeline to generate cortical surface reconstructions and vertex-wise cortical thickness maps [77]. Cortical thickness was averaged across all surface vertices belonging to each of the 400 regions, ensuring topologically accurate regional summaries while preserving anatomical accuracy.

#### Cerebral Blood Flow

To quantify cerebral blood flow, whole™brain CBF maps were calculated from perfusion weighted images [78]. Image processing included motion, distortion and partial volume correction, a macro vascular component, adaptive spatial regularisation of perfusion and T1 uncertainty. The CBF images were aligned to the anatomical T1 images and normalised to MNI152 space and values calculated for the Schaefer 400 regions for each participant.

#### PET Processing and Brain Feature Extraction

The list-mode PET data were binned into 344 3D sinogram frames of 16s intervals, corrected for attenuation and reconstructed using Ordinary Poisson-Ordered Subset Expectation Maximization algorithm (3 iterations, 21 subsets) with point spread function correction. The reconstructed DICOM slices were converted to NIFTI format with size 344 × 344 × 127 (size: 1.39 × 1.39 × 2.03 mm^3^) for each volume. We formed a single 4D NIFTI volume by concatenating the 3D volumes.

The PET volumes were motion [79] and partial volume corrected [77], applying a 25% grey matter threshold [80]. The PET images were also spatially smoothed [81] using a Gaussian kernel with a full width at half maximum of 8mm and co-registered to MNI space.

##### Metabolic connectivity

Metabolic connectivity matrices were derived from dynamic fPET data for use in metabolic gradient analysis and calculation of degree centrality. For each subject, connectivity matrices were constructed using Pearson correlation between the 400 regions. The timeseries included frames 75-188, corresponding to the steady-state FDG uptake period [19, 33].

##### Degree

To calculate fPET degree, the 400 x 400 region correlation matrices were thresholded using a density-based approach that preserved the top 10% of strongest connections, ensuring consistent network sparsity across subjects. From these thresholded adjacency matrices, we computed degree centrality for each region as the sum of its suprathreshold connections, providing a measure of regional integration within the whole-brain metabolic network.

##### Metabolic rate of glucose

Calculations of CMR_GLC_ were undertaken using the FDG time activity curves for the Schaefer atlas 400 regions. The FDG in the plasma samples was decay-corrected for the time between sampling and counting as the input function to Patlak models [82].

##### Glucodynamic variability and complexity

We calculated *sample entropy* and the *standard deviation* of the fPET timeseries for each participant. Entropy was calculated per voxel using embedding dimension m=2 and tolerance r=0.15 [31, 33]. The standard deviation was also calculated voxel-wise to capture variability. Regional entropy and standard deviation values were obtained by averaging voxel-wise values within each of the Schaefer 400 atlas regions.

### Gene Expression

Regional gene expression data were derived from microarray measurements of six post-mortem brains provided by the Allen Human Brain Atlas [36]. The gene data was processed using the Abagen toolbox [83] to map expression values to the Schaefer 400 regions. Expression values for samples assigned to the same brain region were averaged within each donor and then across donors, resulting in a final regional expression matrix of 400 brain regions by 15,631 genes. From this matrix, we extracted expression profiles for 473 unique genes involved in glucose metabolism, identified from the Gene Ontology (GO:0006006), MSigDB Hallmark Glycolysis, and KEGG Glycolysis and Gluconeogenesis pathways. We analysed the 473 genes against multiple ontology sources using enrichment analyses [39], confirming enrichment for core glucose metabolism pathways including glucose metabolic process, regulation of glucose metabolism, and energy generation pathways.

### Analyses

#### Metabolic Gradient–Brain Features

Participants’ metabolic gradients were computed from their 400 x 400 connectivity matrix using diffusion map embedding [37], reducing the high-dimensional connectivity space into its dominant components. To align gradients across subjects to a common space, we performed iterative Procrustes alignment (n=10 iterations). Four gradients were identified, capturing the dominant axes of metabolic variation across the cerebral cortex. We computed group-level gradient maps and subject-specific gradient loadings at the 400 region level, enabling multi-scale investigation of metabolic brain organisation.

#### Metabolic Gradient–Brain Feature Associations

We assessed the biological basis of the metabolic gradients by correlating gradient loadings with multiple neurobiological measures across the 400 regions. The four gradients were correlated with 10 target variables: baseline rates of glucose metabolism (CMR_GLC_), the standard deviation and entropy of the fPET timeseries (fPET_SD_ and fPET_EN_), cerebral blood flow (CBF), fPET degree, cortical thickness, and four fMRI-derived functional gradients. Pearson correlations were computed for each gradient-target pair, with significance assessed at p-FDR < 0.05.

#### Metabolic Gradients and Gene Expression

To identify multivariate associations between the four metabolic gradients and gene expression, we performed a Partial Least Squares Correlation (PLSC) analysis between 400-region matrices of the gradients and the expression of 473 genes involved in glucose metabolism. The matrices were z-scored and the PLSC computed via a singular value decomposition (SVD) of the cross-covariance matrix, yielding latent variables (LVs) that maximise the covariance between the two datasets. The statistical significance of each LV was assessed using permutation testing (n=1,000). The stability of the gradient and gene contributions to each significant LV was evaluated using bootstrap resampling (n=500). From this, we calculated bootstrap ratios (BSRs) for each feature, thresholding at |BSR| > 2.0 to identify robustly contributing gradients and genes.

To characterise the significant genetic metabolic pathway and process associated with the metabolic gradients, the significant genes from the PLSC were subject to an enrichment analysis [39]. The following ontology sources were used: KEGG Pathway, GO Biological Processes, Reactome Gene Sets, Canonical Pathways, CORUM and PANTHER Pathway. Terms with a p-value < 0.01, a minimum count of 3, and an enrichment factor > 1.5 (ratio between the observed counts and the counts expected by chance) were grouped into clusters based on their membership similarities and tested for significance based on FDR-corrected values.

#### Age-Related Gradient Analysis

We examined age-related differences in the metabolic gradient by comparing younger (N=40) and older adult (N=44) people on four gradient metrics. For each participant, we computed (1) the sum of absolute gradient values, and (2) the standard deviation across 400 regions across the four gradients. To specifically test gradient strength at the poles, we identified the top 10% of parcels (N=40) with largest (3) positive, and (4) negative loadings for each gradient and calculated their mean absolute value. Age group comparisons were performed using two-sample *t*-tests (2-sided).

#### Metabolic Gradients and Cognition

We used principal components and regression analyses to test the relationship between metabolic gradients and cognition. First, principal components was used to reduce dimensionality in the cognitive data. The four cognitive measures were converted to Z-scores and entered in a Principal Component Analysis (PCA) with varimax rotations to identify components with eigenvalues greater than one. Participant component scores were saved for use in further regression analysis. The cognition principal component was predicted from the four gradient measures: sum and standard deviation of gradient loadings and loadings among the top 10% of regions with the largest positive and negative loadings.

#### Outlier detection

Outlier detection was also applied across the pipeline: subjects exceeding ±3 standard deviations on the connectivity matrices or brain feature measures were excluded pairwise from subsequent analyses that included that measure. We report the n-size of each analysis.

## Data and Code Availability Statement

The datasets and code used for the study are available from at https://osf.io/r8p69/overview.

## Author Contributions

SDJ and GFE conceived the project. HD and SDJ, CM and GFE designed and developed the manuscript. HD and EL analysed the data. HD wrote the manuscript, SDJ, GFE and CM reviewed/edited the manuscript. All authors read and approved the final manuscript.

## Acknowledgements

This work was supported by Australian Research Council (ARC) Discovery Project DP25010302 and ARC Fellowship FT250100206.

We thank Robert Di Paolo, Gerard Murray, M. Navyaan Siddiqui, Katharina Voigt, Richard McIntyre, Lauren Hudswell and the staff at Monash Biomedical Imaging for their contributions to data acquisition and image reconstruction.

