## Supplement for "Metabolic Connectivity Gradients of the Human Brain"

**~ Supplementary Information ~**

#### Contents

1. Participants – demographics, anthropometry & cognition ____ 2
2. Cognitive Tests ____ 3
3. fMRI Gradients ____ 4
4. Functional connectivity gradient associations with brain physiology ____ 5

#### This file includes:

Supplementary text
Figs. S1
Tables S1

### 1. Participants – demographics, anthropometry & cognition

The characteristics of the whole sample (N =84), as well as the younger (N = 40) and older (N = 44) participants, are shown in Table S1. The mean age of the whole sample was 53.1 years (SD = 24.9). The proportion of women was 50%. The average years of education was 17. Average BMI was 25.0 $kg/m^2$, resting heart rate was 78 BPM and systolic and diastolic blood pressure were 135 and 82 mmHg, respectively. The mean fasting blood glucose was 5.0 mmol/L.

The mean age of the younger group was 27.8 years and the older group 75.5 years. The proportion of women in the younger group (60%) and older group (48%) was not significantly different. The average years of education was higher in the younger (18.0) than the older group (16.2). These years of education are slightly above the average adult population in Australia [1].

The older group had significantly higher mean systolic blood pressure than the younger group (150mmHg vs 119mmHg). The older group also had a higher fasting blood glucose level (5.2 vs 4.8 mmol/L). The older group also has significantly worse performance on the HVLT (7.1 vs 9.3), category switch reaction time (2.16 vs 1.42s) and digit symbol substitution (27.3 vs 63.6) performance.

**Table S1.** Demographics for the whole sample and comparison of older and younger groups. Continuous variables are mean (standard deviation); categorical variables are number and percentage.

| | Whole Sample (N = 84) | | Younger (N = 40) | | Older (N = 44) | | Young vs Old p-value |
|---|---|---|---|---|---|---|---|
| | Mean | SD | Mean | SD | Mean | SD | |
| Age (years) | 53.1 | 24.9 | 27.9 | 6.2 | 76.0 | 6.0 | NA |
| Sex (% female) | 51.0 | | 50.0 | | 46.0 | | 0.659 |
| Education (years) | 17.0 | 3.6 | 18.0 | 2.7 | 16.1 | 4.0 | 0.020 |
| Fasting blood glucose (mmol/L) | 5.0 | 0.5 | 4.8 | 0.4 | 5.2 | 0.6 | < 0.001 |
| Systolic Blood Pressure (mmHg) | 135.4 | 26.5 | 119.0 | 17.2 | 150.4 | 24.7 | < 0.001 |
| Diastolic Blood Pressure (mmHg) | 81.5 | 12.9 | 78.5 | 13.1 | 84.3 | 12.3 | 0.036 |
| Resting Heart Rate (bpm) | 78.3 | 15.6 | 82.5 | 16.9 | 74.6 | 13.4 | 0.020 |
| Body Mass Index ($kg/m^2$) | 24.9 | 4.1 | 24.1 | 4.6 | 25.5 | 3.5 | 0.064 |
| HVLT: Delayed Recall | 8.2 | 2.7 | 9.3 | 2.5 | 7.1 | 2.6 | < 0.001 |
| Category Switch: RT in Switch Trials (sec) | 1.81 | 0.65 | 1.42 | 0.40 | 2.16 | 0.65 | < 0.001 |
| Digit Symbol Substitution: Correct Count | 44.4 | 23.6 | 63.6 | 16.5 | 27.3 | 13.8 | < 0.001 |
| Stop Signal: RT in Stop Signal Trials | 5.7 | 1.2 | 5.3 | 1.3 | 6.0 | 1.2 | 0.076 |

[1]P-values are based on T-test for continuous (2-sided) and Ch-square for categorical variables. One older participant's digit symbol substitution score was more than 3 standard deviations below the mean, and their data was excluded from analyses of cognition

### 2. Cognitive tests

**Hopkins Verbal Learning Test (HVLT).** The HVLT is a three-trial list learning and free recall task. The learning trials comprised 12 words, four words from each of three semantic categories [2]. Approximately 20–25 minutes after the learning trials, participants completed delayed recall and recognition trials. The delayed recall required free recall of any of the 12 words. The recognition trial comprised 24 words, including the 12 target words and 12 false-positives, six semantically related, and six semantically unrelated. Delayed recall was calculated as the total words recalled.

**Task Switching.** For task switching, a computerised test was used in which participants were presented with a word and had to perform a categorisation task. The categorisation task was dependant on two cues that appeared on screen across the trials. One cue was a heart symbol, for which participants were asked to categorise the word presented via a key press as either a LIVING or a NON-LIVING object. If the cue was an arrow-cross, participants were asked to categorise the word as either BIGGER or SMALLER than a basketball. The cue was randomised across trials. Half the trials were switch trials and half were non-switch trials. Half the switch and non-switch trials was congruent in the key presses for either task, half was incongruent. The task switching measure was latency of correctly responding to a switch trial [3].

**Stop Signal.** The stop signal trial was a computer-based test [4] in which participants were required to press the left response key if an arrow on screen pointed left and the right response key if the arrow pointed right. If a signal beep sounded, participants were instructed to stop their response. The delay between presentation of an arrow and signal beep started at 250ms and was altered up or down by 50ms based on performance. The delay increased up to 1150ms if the previous stop signal trial was successful and decreased to 50ms if the previous trial was unsuccessful. The stimulus onset asynchrony between the onset of a fixation circle at the start of each trial was 2000ms. Reaction time in the stop signal trials was recorded.

**Digit Symbol Substitution.** A computer-based task presenting participants with a matrix of 18 column and 16 rows [5]. Participants were required to translate symbols shown above the matrix in a key into digits in the matrix. The trial lasted two minutes. Performance was measured as total count of correct responses.

#### 3. fMRI gradients

Four fMRI gradients were identified explaining the following variance: $FC_{G1}$=36.7%, $FC_{G2}$=19.7%, $FC_{G3}$=31.5%, and $FC_{G4}$=12.0% (see Figure 2, main manuscript). Gradient 1 represents a primary sensory-to-association hierarchy. The positive pole reflects heteromodal association areas, featuring regions from the default, salience ventral attention, and control networks. In contrast, the negative pole is strongly anchored in unimodal sensory regions, including primary and peripheral visual areas and primary somatomotor cortices, indicating a fundamental axis from concrete sensory processing to abstract, transmodal,.

$FC_{G2}$ captures a functional differentiation between the dorsal attention and somatomotor and visual systems. The positive pole is dominated by regions of the dorsal attention network, including key nodes in the superior parietal lobule and temporo-occipital cortex, alongside somatomotor and auditory regions. The negative pole comprises transmodal and limbic areas, including temporal pole, prefrontal, and posterior cingulate regions, suggesting an axis that separates externally-oriented, spatially-focused attention from internal processing.

$FC_{G3}$ delineates the default network and temporal association cortex from the somatomotor cortex. The positive pole is characterized by widespread regions of the default mode and temporal parietal networks, including temporal prefrontal, and posterior cingulate and retrosplenial areas. The negative pole includes visual and somatomotor regions, positioning this gradient as one that primarily differentiates the brain's intrinsic, self-referential system from primary sensorimotor cortices.

$FC_{G4}$ describes a clear somatomotor-dorsal attention versus visual processing segregation. The positive pole is characterised by somatomotor and dorsal attention networks. The negative pole is almost exclusively composed of visual regions from both central and peripheral areas. This gradient highlights a fundamental organizational principle that distinguishes body-centred sensation and action from exteroceptive visual perception.

### 4. Functional connectivity gradient associations with brain physiology

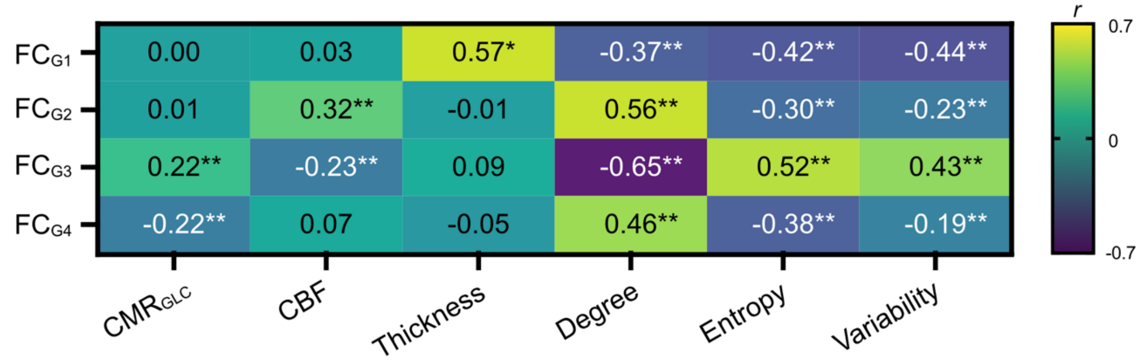


**Figure S1. Functional gradient and neurophysiological brain feature associations.** Correlation of metabolic gradients loadings with brain features. *p-FDR < 0.05; **p-FDR < .001.

The functional connectivity gradients demonstrated select significant associations with the neurobiological measures (Figure S1), with fewer overall significant associations than was found for the metabolic connectivity gradients. Unlike the primary metabolic connectivity gradient ($MC_{G1}$), $FC_{G1}$ was not significantly associated $CMR_{GLC}$ nor CBF (r = 0.00 and 0.03, p > .05). $FC_{G2}$, $FC_{G3}$ and $FC_{G4}$ showed varied positive and negative associations with $CMR_{GLC}$ and CBF. $FC_{G3}$ positively associated with $CMR_{GLC}$, (r = 0.22 p-FDR < 0.001) and negatively associated with CBF (r = -.23, p < .001), indicating that regions of the default mode and temporal parietal networks at the positive gradient pole had higher $CMR_{GLC}$ but lower CBF than the visual and somatomotor regions at the negative gradient pole. $FC_{G4}$ was also negatively associated with $CMR_{GLC}$, indicating that somatomotor and dorsal attention networks at the positive pole of the gradient had lower $CMR_{GLC}$ than visual regions from both central and peripheral areas at the negative pole of the gradient.

Cortical thickness was positively associated with $FC_{G1}$ (r = 0.51, p < .001). The positive pole including regions from the default, salience ventral attention, and control networks had greater cortical thickness than the visual and somatomotor regions at the negative pole. Unlike the metabolic connectivity $MC_{G2}$ and $MC_{G3}$, $FC_{G2}$ and $FC_{G4}$ were not significantly associated with cortical thickness.

Functional network degree showed varied patterns of positive and negative associations with the functional gradients. $FC_{G1}$ and $FC_{G3}$ were negatively associated with functional network degree (r = -0.37 and r = -0.65, p < .001), whereas $FC_{G2}$ and $FC_{G4}$ were positively associated with functional network degree (r = 0.56 and r = 0.46, p < .001). These associations were in the opposite direction to those observed for the metabolic connectivity gradients ($MC_{G2}$ and $MC_{G4}$), indicating that the transmodal regions ($FC_{G1}$) and default mode and temporal parietal networks ($FC_{G3}$) had lower functional network degree than there opposing visual and primary somatomotor areas. In contrast, the positive pole regions in the dorsal attention and somatomotor and auditory regions ($FC_{G2}$ and $FC_{G4}$) had higher network degree than transmodal and limbic areas ($FC_{G2}$) and visual areas ($FC_{G4}$).

The functional connectivity gradients showed mostly negative associations with the haemodynamic measures, with the exception of $FC_{G3}$. Higher entropy (complexity) and complexity (SD) of the fMRI timeseries was significantly associated with lower $FC_{G1}$, $FC_{G2}$ and $FC_{G4}$ expression (r = -0.30 to -0.42, p-FDR < .001; r = -0.23 to -0.44, p < .001). Higher variability and complexity of the fMRI timeseries was significantly associated with higher $FC_{G3}$ expression (r = 0.52 and r = 0.43, p-FDR < .001). For example, for $MC_{G1}$, the positive (prefrontal regions of the control, default, salience ventral and temporal parietal networks) had lower entropy and variability than the negative pole (visual and somatomotor regions). Together, these findings indicate systematic network-level differences in haemodynamics that map onto the brain's functional hierarchy. These results are also consistent with the metabolic connectivity gradient results reported in the main manuscript. Transmodal and executive regions exhibit lower haemodynamic complexity and variability, consistent with flexible network transitions from a relatively stable metabolic baseline. In contrast, unimodal regions show greater haemodynamic complexity and variability, potentially reflecting haemodynamic sensitivity to fluctuating sensory demands.

Taken together, these findings reveal that the functional network organisation of the brain are less strongly associated with $CMR_{GLC}$, CBF and cortical thickness and express different patterns of network connectivity than metabolic connectivity gradients. In contrast, glucodynamic and haemodynamic complexity and variability more consistently map onto their respectively hierarchies.
